# An “alert state” ribosome population acts as a master regulator of cytokine-mediated processes

**DOI:** 10.1101/2023.10.17.562425

**Authors:** Anna Dopler, Ferhat Alkan, Yuval Malka, Kelly Hoefakker, Olga I. Isaeva, Mandy Kerkhoff, Anastasia Gangaev, Joana Silva, Sofia Ramalho, Liesbeth Hoekman, Maarten Altelaar, Rob van der Kammen, Jonathan Wilson Yewdell, Pia Kvistborg, William James Faller

## Abstract

Inflammatory cytokines are pivotal to immune responses. Upon cytokine exposure, cells enter an “alert-state” that enhances their visibility to the immune system. Here, we identified an “alert-state” subpopulation of ribosomes (ASRs) defined by the presence of the P-stalk. We show that ASRs are formed in response to cytokines linked to tumor immunity, and are involved in the preferential translation of mRNAs vital for the cytokine response.

Mechanistically, ASRs are required for the efficient translation of transmembrane domains of receptor molecules involved in cytokine-mediated processes. Importantly, loss of the ASR prevents CD8+ T cell recognition and killing, and inhibitory cytokines like TGFβ hinder ASR formation, suggesting that the ASR is a central regulatory hub upon which multiple signals converge. Thus, the ASR is an essential mediator of the cellular rewiring that occurs following cytokine exposure, via the translational regulation of this process.

## Introduction

Immune evasion is a well-known occurrence in cancer cells, and has been included as one of the new Hallmarks of Cancer^1^. The success of immune checkpoint blockade (ICB) in several cancer types has highlighted the importance of immune surveillance in tumor control, however the majority of patients either do not respond, or acquire resistance to such therapies^2^. It is well established that the infiltration of T cells in the tumor microenvironment (TME) is a critical mediator of ICB^3^. While T cells directly kill tumor cells, they also secrete cytokines such as interferon-γ (IFNγ) and TNFɑ, and as a result, human tumors often show evidence of transcriptional signatures of IFNγ signaling^4^. While IFNγ signaling serves as a useful biomarker of ICB response^5^, it also leads to a major rewiring of tumor cell and immune cell behavior^6^. Upon cytokine exposure, cells take on an “alert state”, that is characterized by an up-regulation of chemokines and checkpoints, an increase in antigen processing and presentation (APP)^7^, and in some cases, the onset of senescence or apoptosis^8^. Conversely, inhibitory cytokines like TGFβ do the opposite, as can be seen in tumors with a dominant immunosuppressive myeloid cell population^9^.

However, despite this major rewiring and the significant phenotypic consequence of it, our understanding of how cytokines modify tumor behavior is still lacking. Analysis of cytokine responses have tended to focus on transcriptional regulation, and while some examples of post-transcriptional regulation of cytokine responses exist^10–12^, there is still much to learn in this context.

The concept of ribosomal populations with specific functions has been around for more than 20 years^13,14^, but it remains controversial. Recent work has clearly shown more variability to the ribosome than has previously been appreciated^15,16^, and this supports the potential of specialized functions^17–20^. Many members of the translation machinery are selectively regulated^21^, and considering its complexity, it is not difficult to imagine the ribosome itself being similar. The stoichiometry of ribosomal proteins (RPs) has been of particular interest in this context^16,22^, and RPs are known to be important for T cell survival and development^23–25^, dendritic cell activation^26^, and B cell activation^27^, demonstrating that they play a role in immune responses. Indeed, the “immunoribosome” is a hypothesized subset of ribosomes that selectively generates antigenic peptides^28^. Recent evidence has supported a role for RPs in this context^29^, suggesting that ribosomes may play an unappreciated role in responses to cytokine exposure.

Based on this, we speculated the existence of a post-transcriptional mechanism of cytokine-induced cellular rewiring, mediated by a change in the ribosome. A cytokine-responsive ribosome would make sense from an evolutionary standpoint, as transcriptional changes are relatively slow, so changes to the ribosome would allow for a more responsive fine-tuning of the “alert state”. Such a regulator of cytokine responses would be of critical importance for our understanding of immune regulation in patient response to ICB.

Here, we identify an cytokine-responsive “alert state” ribosome (ASR) population, defined by the presence of the P-stalk, and show that the ASR is a critical mediator of the cellular rewiring that occurs following cytokine exposure, via the translational regulation of this process.

## Results

### The P-stalk defines a cytokine-responsive “alert state” ribosome (ASR)

To identify RPs that regulate tumor cell response to cytokines, we treated human melanoma cells (Mel624 and M026 cell lines) with IFNγ and TNFɑ. We then performed proteomic analysis of actively translating ribosomes to identify RPs that change in response to treatment (Fig. 1a). We confirmed that the cells took on a cytokine-induced “alert state” by measuring both APP activation by increased HLA I cell surface expression (determined via flow cytometry), and the expression of immunoproteasome subunits (determined by proteomics) (Supp. Fig. 1a-d). We then used sucrose density gradient centrifugation to isolate the actively translating ribosomes from these cells. We also collected total protein, and compared cytoplasmic RP levels to those in the translating ribosomes using LC-MS. This analysis revealed that in both cell lines, P1 was the protein with the largest increase in the translating ribosomes (Fig. 1b). We confirmed this increase by measuring polysome, sub-polysome, and total P1 levels by immunoblotting, which confirmed that P1 is significantly increased in polysomes (actively translating ribosomes) following IFNγ/TNFɑ exposure.

**Figure 1:**
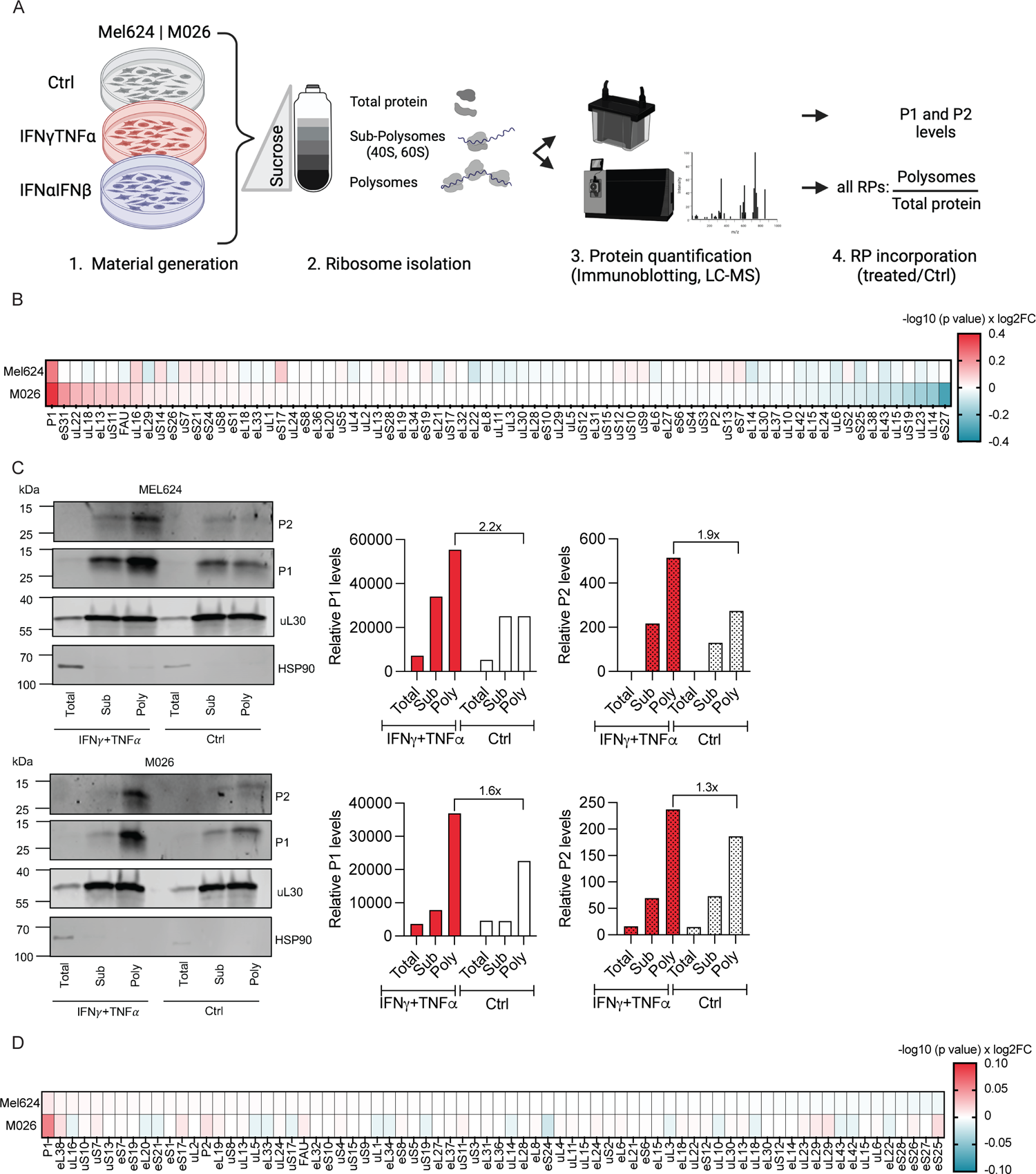
Inflammatory cytokines stimulate ASR formation. **A.** Schematic representation of experimental setup. **B.** P1 association with the translating ribosome (poly) is increased upon IFNγ and TNFɑ exposure in Mel624 and M026 tumor cells. Heatmap shows polysome to total ratio in cytokine-treated versus untreated cells. n=3 independent experiments. **C.** Western blot analysis confirming increased polysomal association of P1 and P2 upon IFNγ and TNFɑ exposure. HSP90 was used as loading control and uL30 was used as control RP for data normalization. n=1 experiment per cell line. **D.** P1 association with the translating ribosome (poly) is increased upon IFNα and IFNβ exposure in Mel624 tumor cells. Waterfall plot shows polysome to total ratio in cytokine-treated versus untreated cells. n=3 independent experiments.

Crucially, another RP, uL30, does not increase, demonstrating that this is not a general effect on the ribosome (Fig. 1c). P1 forms a heterodimer with P2, and two of these heterodimers are anchored to the ribosome via uL10. This pentameric stalk (also known as the P-stalk) is the central component of the GTPase-associated center of the ribosome^30^. Interestingly, the P1/P2 heterodimer has long been known not to be essential for translation^31^, and, unlike most other RPs, dynamically associates with the ribosome^32^. To understand whether the cytokine-mediated changes we have detected are P1 specific, or rather a result in an increase in the P-stalk, we tested whether P2 is also increased in translating ribosomes following cytokine treatment. Despite not identifying it in our proteomic analysis, western blotting showed that this is indeed the case (Fig. 1c), suggesting that the addition of the P1/P2 heterodimer to the ribosome is increased following IFNγ/TNFα exposure.

IFNγ and TNFα are just two of the cytokines known to regulate the “alert state”. Additional experiments revealed that IFNα and IFNβ also increase ASR abundance (Fig. 1d), suggesting that the ASR is activated regardless of the activating signal. We confirmed these findings in cell lines from diverse origins (ovarian cancer (SK-OV-3), colorectal cancer (HCT 116), and immortalized retinal pigment epithelial cells (hTERT-RPE1) (Supp. Fig. 1e-g), demonstrating that a change in ribosomes in response to cytokines is consistent across tissues. These results suggest that the ASR may be a regulatory node upon which multiple signals converge, and that it may therefore act as a general regulator of cytokine-mediated processes.

### The ASR preferentially translates mRNAs important for immunosurveillance

To measure whether the ASR is important for the translation of mRNAs involved in the cytokine response, we used shRNA to knockdown (KD) P1 in both melanoma cell lines, using independent shRNAs (shP1#1, shP1#3). We also used an shRNA against a control RP that was unresponsive to cytokine treatment (shL28), and two scrambled control shRNAs (shCtrl#1, shCtrl#2). Transduction efficiency was measured using mCherry (%) expression and was approximately 90% across cell lines (Supp Fig. 2a). KD efficiency was variable across hairpins and cell lines, with the Mel624 cell line showing a 42% and 50% KD for shP1#1 and shP1#3, and the M026 cell lines showing 28% and 16% KD, respectively (Supp Fig. 2b,c). Interestingly, the strength of the phenotypes presented in the remainder of this manuscript showed a dose dependence, with the Mel624 cell line regularly providing more reliable results, particularly compared to the M026 shP1#3 cell line (16% KD).

**Figure 2:**
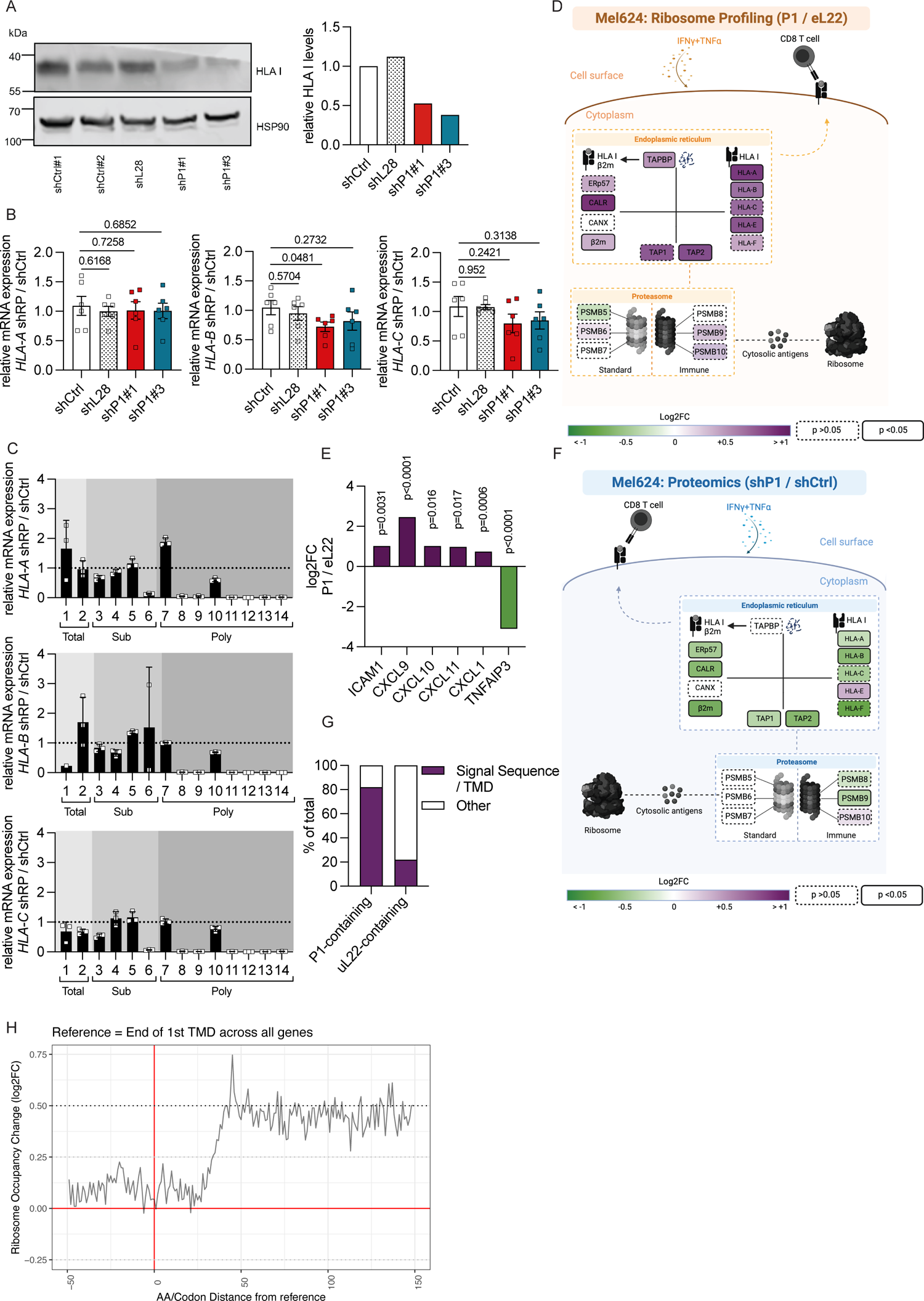
The ASR preferentially translates mRNAs important for immunosurveillance. **A.** Western blot showing decreased HLA I protein levels in shP1 tumor cells. HSP90 was used as loading control. n=1 independent experiment. **B.** RT-qPCR showing no change in HLA mRNA expression (*HLA-A, HLA-B, HLA-C*) following P1 KD using *ACTB* as a reference. n=2 independent experiments each assessed in technical triplicates. Data are represented as mean ± SEM and p values were determined using a two-tailed t-test. **C.** RT-qPCR of isolated fraction 1-14 showing decreased polysomal association of HLA mRNA in shP1 cells, using *ACTB* as a reference. n=3 technical replicates represented as mean ± SD. **D.** Ribosome profiling reveals preferential translation of HLA and other APP components by the ASR. n=2 biological replicates per condition and p values were calculated using the DESEQ2 package. **E.** The ASR preferentially translates key chemokines and other cytokine-responsive mRNAs important for immune response. n=2 biological replicates per condition and p values were calculated using the DESEQ2 package. **F.** Proteomics analysis reveals down-regulation of HLA and other APP components upon P1 KD. n=3 independent experiments and p values were calculated using a two-tailed t-test. **G.** mRNAs that are preferentially translated by ASR are enriched for coding signal peptide and/or transmembrane domains. **H.** The ASR is specifically enriched at translating mRNA regions that come after the translation of the first transmembrane domain. The plot shows the aggregated ribosome occupancy differences (ASR vs all ribosomes) at each codon distance from the reference set, ends of the first TMD domain across all genes.

Following P1 KD, we measured the intracellular protein levels of HLA I (a well-described IFNγ-regulated gene) using a pan HLA-A, -B, -C specific antibody. This showed that total HLA I protein levels were significantly decreased (Fig. 2a, Supp. Fig. 2d). However, qPCR analysis did not reveal changes in mRNA levels (Fig. 2b, Supp. Fig. 2e), suggesting post-transcriptional regulation. Isolation of ribosomal fractions from these cells using sucrose gradient density centrifugation showed that despite no change in HLA-A, -B, or -C mRNA levels, their polysomal association was significantly decreased following P1 deletion, showing that it is translationally regulated (Fig. 2c).

To gain a more global view of the translatome of the ASR, we expressed a HA-tagged version of P1 in the Mel624 cell line, alongside a line expressing HA-tagged eL22 (Supp. Fig. 3a). This allowed us to specifically isolate P1- and eL22-containing ribosomes following IFNγ/TNFα exposure, and characterize the associated mRNAs via ribosome profiling.

Immunoblotting confirmed enrichment of the HA tag following immunoprecipitation (Supp. Fig. 3b), while analysis of the resulting ribosome profiling confirmed a 29 - 30 nucleotide read length and 3-nucleotide periodicity in these data (Supp. Fig. 3c-f). These data showed that alongside HLA I, the mRNAs of many other components of the APP machinery were preferentially associated with the ASR, including β2M, TAP1, TAP2, TAPBP, and CALR (Fig. 2d). Additionally, key chemokines (including CXCL1, 9, 10 and 11), were enriched on the ASR, as were other cytokine-regulated mRNAs including a central regulator of T cell recruitment (ICAM1)^33^, while a negative regulator of TNFα-mediated apoptosis (TNFAIP3)^34^ was depleted (Fig. 2e).

Extending these findings, we carried out a proteomic analysis of shP1 cells compared to shCtrl cells. These data also showed that the protein levels of multiple APP components were decreased following P1 KD (Fig. 2f, Supp. Fig. 3g). Furthermore, as we had seen in the ribosome profiling data, GSEA showed that genesets associated with antigen presentation and HLA I were among the most down-regulated following P1 KD (Supp. Fig. 3h), confirming that multiple members of this process are regulated in the same manner as HLA I.

In order to understand the mechanism by which the ASR regulates specific mRNAs, we analyzed the ribosome profiling data in more detail. Annotation of the sequence features enriched on mRNAs associated with the ASR revealed that messages translated at the ER were substantially enriched. While 22% of eL22-enriched mRNAs contained a signal sequence or a transmembrane domain (TMD), 82% of P1-enriched messages contained them (Fig. 2g). Visualization of the distribution of sequencing reads on TMD containing mRNAs revealed that P1-containing ribosomes were significantly enriched after the first TMD (Fig. 2h), suggesting that the role of the P-stalk in elongation may be causing differential translation of specific mRNAs^35,36^. It has previously been reported that the P-stalk is required for dengue viral replication^37^, as it appears to be needed to relieve ribosome pausing on the viral RNA^38^. The same study suggested that the P-stalk aids in TMD translation, due to pauses that are thought to occur on such sequences^38^, potentially explaining the observed findings.

### The ASR regulates cytokine-mediated processes

In order to assess the functional relevance of translational regulation by the ASR, we assessed several well defined processes. We initially measured whether the ASR-mediated intracellular down-regulation of HLA I reduced APP in cells without cytokine exposure. This was indeed the case, and loss of the ASR resulted in a strong and significant decrease of both pan class I and HLA-A2 cell surface expression compared to cells infected with shL28 or a scrambled control (Fig. 3a; Supp. Fig. 4a). We also measured the ability of these cells to recover their surface HLA I after acid-mediated removal, and found that while the kinetics of recovery were similar, the magnitude was significantly reduced following P1 KD, as would be expected in cells with reduced HLA I levels (Fig. 3b; Supp. Fig. 4b). We then treated the cells with IFNγ/TNFα, and measured their ability to take on the “alert state”, which we measured via the up-regulation of HLA I cell surface expression. While all cells showed an increase in cell surface HLA I, that increase is significantly attenuated in shP1 cells compared to shL28 or scrambled controls (Fig. 3c; Supp. Fig. 4c).

**Figure 3:**
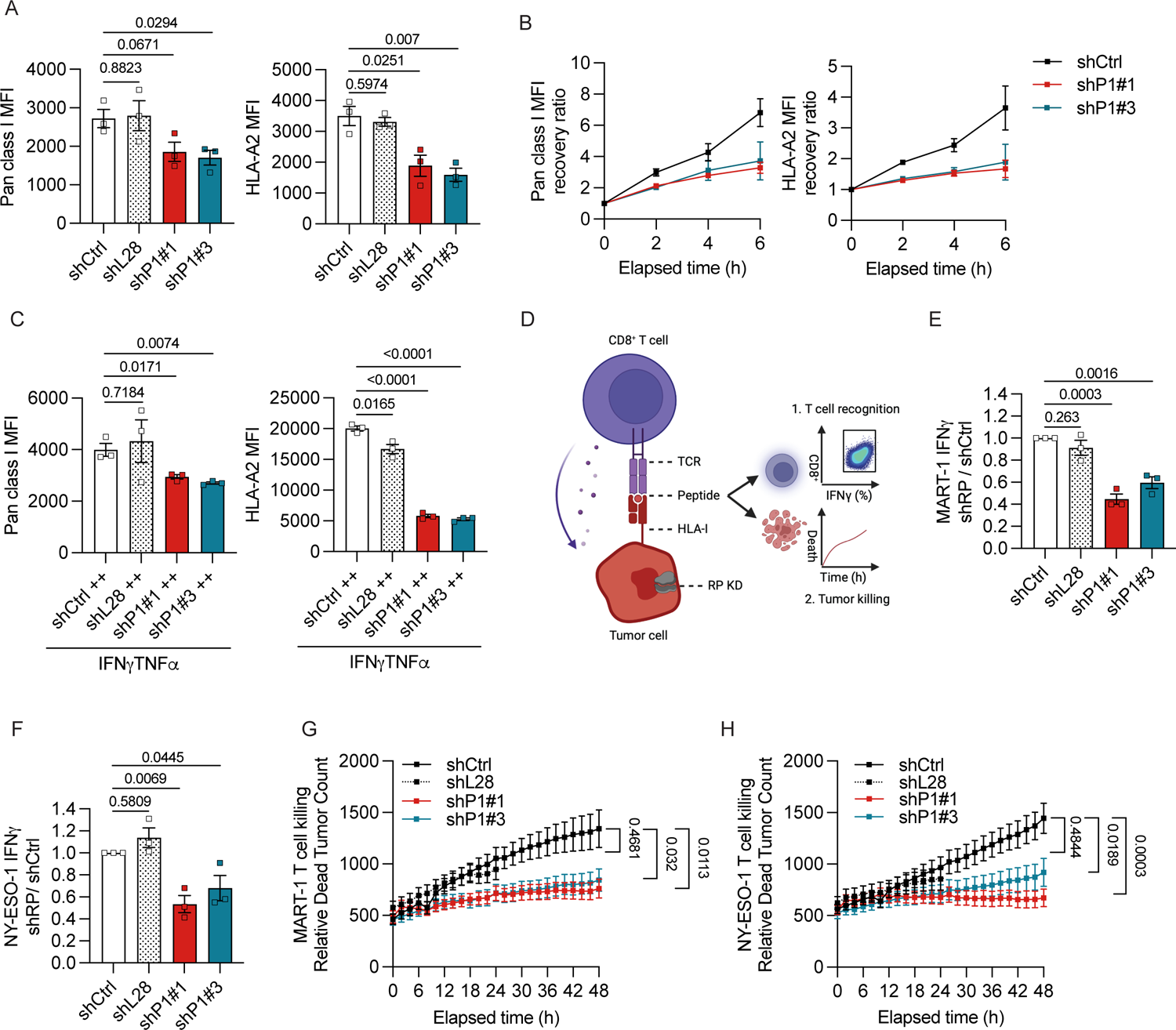
The ASR regulates cytokine-mediated processes. **A.** P1 KD decreases HLA I surface levels. HLA levels were assessed by flow cytometry using the MFI. n=3 independent experiments each assessed in triplicates. Data are represented as mean ± SEM and p values were determined using a two-tailed t test. **B.** shP1 tumor cells showing slower recovery kinetics of HLA I after acid wash compared to the control. n=3 independent experiments. Data are represented as mean ± SEM. **C.** Exposure to IFNγ and TNFα results in low basal HLA I surface levels in shP1 compared to control tumor cells. n=3 biological replicates represented as mean ± SEM. p values were calculated using a two-tailed t-test. **D.** Schematic representation of co-culture experimental set up. **E.** MART-1-specific CD8^+^ T cell recognition is decreased in shP1 compared to control in Mel624 tumor cells. n=3 independent experiments each assessed in triplicates. Data are represented as mean ± SEM and p values relative to shCtrl were determined using a two-tailed t-test. **F.** NY-ESO-1-specific CD8^+^ T cell recognition is decreased in shP1 compared to in Mel624 control tumor cells. n=3 independent experiments each assessed in triplicates. Data are represented as mean ± SEM and p values relative to shCtrl were determined using a two-tailed t-test. **G.** MART-1 specific T cell killing is suppressed in shP1 tumor cells. n=3 independent experiments per cell line, each assessed in triplicates. Data represent mean and error ± SEM and p values were calculated with a two-tailed t-test. **H.** NY-ESO-1 specific T cell killing is suppressed in shP1 tumor cells. n=3 independent experiments per cell line, each assessed in triplicates. Data represent mean and error ± SEM and p values were calculated with a two-tailed t-test.

We then carried out a melanoma cell/T cell co-culture assay^39^. This assay is based on donor CD8+ T cells retrovirally transduced with a TCR specific for HLA-A2 restricted peptides derived from MART-1 or NY-ESO-1, both of which are presented by Mel624 and M026 tumor cells (Fig. 3d). If antigens are correctly presented, they will be recognized by T cells, resulting in their activation (measured by IFNγ and TNFα production, and CD107a cell surface expression), and the killing of the tumor cells. While KD of eL28 did not affect T cell recognition, KD of P1 resulted in a strong and significant decrease in recognition by both MART-1 and NY-ESO-1 specific CD8+ T cells (Fig. 3e,f; Supp. Fig. 4d,e). A live imaging killing assay confirmed this finding, with shP1-expressing cells being far less efficiently killed by T cells compared to either shL28 or a scrambled control (Fig. 3g,h; Supp. Fig. 4f,g).

As KD of P1 may affect the ribosome more generally, it is important to ensure that the observed effect is not a result of reduced expression of the source proteins from which the antigens derive. However, the levels of MART-1 and NY-ESO-1 were largely unchanged in shP1 cells (Supp. Fig. 5a), discounting this possibility, although P1 KD did reduce both protein synthesis (Supp Fig. 5b) and proliferation (Supp Fig. 5c). We next inhibited protein synthesis to the same degree by knocking down another RP, eL6 (Supp. Fig. 5d). This intervention did not cause a decrease in HLA I cell surface expression, consistent with the conclusion that the effect of the ASR is not based on its effect on global protein synthesis (Supp. Fig. 5e). Additionally, as the P-stalk has been shown to regulate GCN2 activation^40^, we also measured the levels of phosphorylated elF2α in both cell lines. These data showed no difference in elF2α phosphorylation following P1 KD, either with or without IFNγ/TNFα treatment (Supp Fig. 5f) suggesting that this role of the P-stalk is not relevant following cytokine exposure.

To explore the clinical relevance of our findings, we turned to the TCGA database^41^. RPs tend to be co-regulated, so we therefore ranked samples based on P1 or P2 expression relative to the expression of other RPs, thus controlling for globally high/low RP expression, and enabling us to focus on samples whose P1 or P2 levels are higher than other RPs. This analysis showed that high P1 or P2 levels (relative to other RPs) significantly positively correlated with an IFNγ signature^42^ and a effector CD8 T cell signature^4^ in melanoma (Supp Fig. 5g, h). Crucially, we did not observe the same correlation when other RPs were analyzed in the same manner (Supp Fig. 5g, h). Further analysis of TCGA data revealed the same significant correlation in the majority of cancer types (24 of 36), suggesting that the P-stalk regulates cytokine responses in diverse tissues (Fig. 4a).

**Figure 4:**
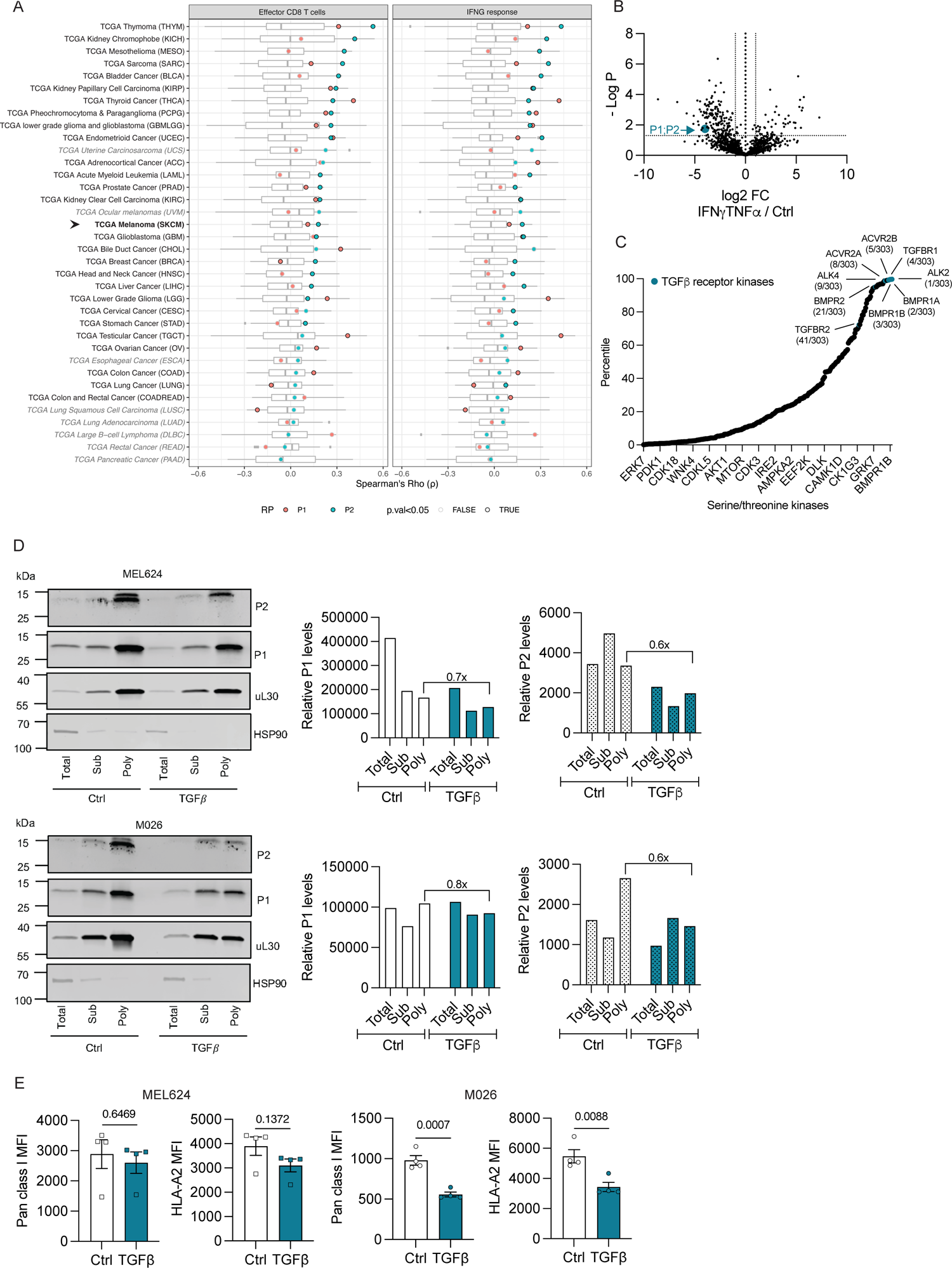
TGFβ inhibits ASR formation. **A.** Correlation analyses of TGCA cohorts (n=36) and all RPs (n=90) reveals a significant correlation between relative P-stalk expression (z-score) and IFNγ response signature and effector CD8 T cell infiltration in diverse tissues. Box plots represent the distribution of RP-specific correlation coefficients (Spearman’s rho). Significant correlations (p<0.05) are circled in black. **B.** IFNγ/TNFα exposure results in loss of P1/P2 phosphorylation. n=3 independent experiments**. C.** Prediction of P1 and P2 phosphorylation by TGFβ receptor kinases (top 5 predicted). **D.** TGFβ inhibits formation of the ASR. Western blot analysis showing decreased polysomal association of P1 and P2 following TGFβ exposure. HSP90 was used as loading control and uL30 was used as control RP where no changes could be observed. n=1 experiment per cell line. **E.** TGFβ decreases HLA I surface levels in Mel624 and M026 tumor cells. HLA levels were assessed by flow cytometry using the MFI. n=4 biological replicates. Data are represented as mean ± SEM and p values were determined using a two-tailed t test.

In all, these findings indicate that the loss of the ASR has a striking effect on the antigen presenting ability of cells, and thus the ability of CD8+ T cells to detect and kill them.

### TGFβ inhibits ASR formation

The P-stalk proteins are known to be phosphorylated, and while the function of this modification on P1 and P2 is unknown, it has been suggested that phosphorylation of the third P-stalk protein (uL10) regulates its association with the ribosome^43,44^. As the phosphorylation sites are conserved among all three P-stalk members^43^, it is reasonable to consider that modification of P1 and P2 may also be regulating their ribosome association. We therefore carried out a phospho-proteomic analysis of the M026 cell line, with and without IFNγ/TNFα. These data showed that there is a loss of phosphorylation upon treatment, suggesting that this is an inhibitory modification (Fig. 4b).

Several kinases have been shown to phosphorylate P1 and P2^43,45^. However, to get a more global view of the potential P1/P2 kinases, we used the recently published serine/threonine kinome^46^. The top 5 kinases predicted to phosphorylate P1 and P2 were all TGFβ receptor kinases (Fig. 4c). TGFβ is a well described inhibitory cytokine, so repression of the ASR via the phosphorylation of P1/P2 would be in keeping with this function. We therefore treated both cell lines with TGFβ, and measured P1 and P2 incorporation into the ribosome. This showed that P-stalk incorporation is decreased by TGFβ (Fig. 4d). Furthermore, HLA I cell surface expression was down-regulated, supporting the finding that TGFβ inhibits the ASR (Fig. 4e).

## Discussion

It is well-established that for the majority of cancers, the expression of chemokines and HLA I is strongly correlated with clinical response to checkpoint targeting therapies^47–49^. Indeed, many alterations that affect the “visibility” of a tumor are known to be mechanisms of cancer immune evasion, and the silencing of both chemokines and components of the APP machinery have been observed in human cancer^50–52^. As such, the discovery of a novel regulator of these genes is an important one.

Here, we show that: first, the ASR is regulated by both pro- and anti-inflammatory cytokines, positioning it as a central nexus, upon which several signals converge; second, the ASR selectively translates many cytokine-responsive mRNAs, including APP components, chemokines, and other mRNAs that help rewire cells following cytokine exposure; third, this may be regulated by a phosphorylation/dephosphorylation cycle; and fourth, that this is a general mechanism across tissue and cancer types.

The fact that the ribosome itself is at the center of this regulation is in fitting with recent reports of direct regulation of phenotypes by the ribosome^22,53–55^. Indeed, it has previously been hypothesized that an “immunoribosome” exists^28^, which was defined as a subset of ribosomes dedicated to translating DRiPs for HLA I immunosurveillance^13,14^, thus altering the antigen repertoire. However, the ASR plays a much broader role than this, translating the mRNA involved in multiple cytokine responses, and acting as an “alert state” ribosome, regulating not just APP, but much of the cellular rewiring that occurs upon cytokine exposure. While the ribosome has not traditionally been considered a viable target for therapy due to its crucial role in biology, the recent identification of dynamic RP/ribosome interactions suggest that there may in fact be unrealised opportunities^54,56^. Indeed, the P1/P2 stalk proteins were the first RPs shown to dynamically associate with ribosomes^57^, highlighting them as interesting RPs in this context.

It remains to be fully determined why and how the ASR preferentially translates specific mRNAs. Interestingly, it has been described previously that the ribosomal P-stalk is needed for the translation of transcripts encoding TMDs^38^, and we see a similar effect here. As a result of their function, cytokine responsive proteins are enriched for signal peptides and TMDs, such as HLA I and the TAP transporter proteins, which could at least partially explain the results described here. However, many signal peptide and TMD containing mRNAs are not enriched on the ASR, suggesting that this is only part of the picture. Other potential explanations could include mRNA selection via the binding of specific RNA binding proteins to mRNAs^58^, or altered ribosome localization. An attractive hypothesis that links the ASR to the original concept of the immunoribosome, is that the P-stalk localizes a subset of ribosomes to TAP-associated proteasomes to enable peptide channeling^59^, but such additional mechanisms were not probed in this study.

Our discovery also raises several additional questions. We have shown that P-stalk incorporation into the ribosome is regulated by both inflammatory (IFNγ/TNFα, and IFNα/IFNβ) and inhibitory (TGFβ) cytokines, but we have not extensively tested the ability of all relevant cytokines to do the same. It will also be interesting to know how widespread this role of the ASR is. Our analysis of TCGA datasets suggest that cytokine response is regulated in this manner in a large number of tissues. It would be particularly interesting to extend studies of the ASR to professional antigen presenting cells like dendritic cells, where RPs are known to be highly regulated during activation^26^.

In the context of cancer, the down-regulation of P-stalk proteins or the creation of the ASR could plausibly be a mechanism through which tumor cells avoid immunosurveillance, or resist immunotherapy. We have shown that P-stalk mRNA correlates with IFNγ and CD8+ T cell signatures, however, these data are from bulk RNA Seq, and we cannot prove that tumor P-stalk levels are causing this, rather than P-stalk levels in other contributing cell types. Additionally, as expression levels are not necessarily indicative of P-stalk incorporation into the ribosome, this is likely to be a sub-optimal measure of ASR abundance. Regardless, we believe that the data we have described here shows that further analysis of patient data is warranted.

To summarize, we have identified a novel master regulator of cytokine responses that acts at the level of translational control of gene expression. While the technologies to study specialized ribosomes in patient tissue in a high-throughput manner are currently lacking, it will be an important feature to consider in future studies, and may present therapeutic applications.

## Supplemental figure legends

## Methods

### Patient material

Human peripheral blood mononuclear cells (PBMCs) were derived from buffy coats of donors at the Netherlands Cancer Institute (Amsterdam, The Netherlands). PBMC isolation was performed using a standard Ficoll gradient centrifugation separation. Isolated PBMCs were either processed directly or cryopreserved in liquid nitrogen and fetal bovine serum (FBS) with 10% dimethyl sulfoxide (DMSO).

### Generation of MART-1 and NY-ESO-1 TCR CD8 T cells

TCR transduced CD8^+^ T cells were isolated from PBMCs as described previously^60^. After PBMC isolation, CD8^+^ T cells were isolated with the Miltenyi CD8^+^ T Cell Isolation Kit, human (Miltenyi Biotec 130-096-495) and stimulated with Dynabeads Human T-Activator CD3/CD28 (Miltenyi Biotec 111.31D) for 48 hours at 37°C. Activated CD8^+^ T cells were mixed with MART-1 or NY-ESO-1 TCR containing retrovirus, added on a retronectin-coated (Takara T100A) non-tissue culture 24-well plate and spun for 90 minutes at 2500 rpm at 10°C. Subsequently, TCR-transduced CD8^+^ T cells were harvested and maintained in T cell medium containing Roswell Park Memorial Institute 1640 Medium (RPMI) supplemented with 100 U/ml Penicillin-100ug/ml Streptomycin and 10% human serum (HS, Sigma-Aldrich), IL-7 (ImmunoTools, 5ng/ml), IL-15 (ImmunoTools, 5ng/ml). After 7 days, TCR transduction efficiency was measured by flow cytometry using the PE a-mouse TCRβ antibody (clone H57-597, BD Biosciences 553172) and anti-PE MicroBeads (Miltenyi Biotec 130-048-801). Finally, TCR-transduced CD8^+^ T cells were rapidly expanded for 5 days using irradiated feeder PBMCs and medium containing 60U/ml IL-2 (Proleukin, Novartis) and Ultra-LEAF™ Purified anti-human CD3 Antibody (clone OKT3, BioLegend 317326, 1:2000). After 5 days, T cells were expanded for additional 2 weeks in medium without the Ultra-LEAF™ Purified anti-human CD3 Antibody and stored at −80°C until needed.

### Cell culture

M026, Mel624, hTERT-RPE1, SK-OV-3 and HCT 116 cells were maintained in Dulbecco’s Modified Eagle Medium (DMEM) supplemented with 10% FBS and 100 U/ml Penicillin-100ug/ml Streptomycin (Life Technologies 15140-122). HEK293T cells were maintained in Iscove’s Modified Dulbecco’s Medium (IMDM) supplemented with 10% FCS and 100 U/ml Penicillin-100ug/ml Streptomycin.

### Cytokine treatment

Cytokine stimulation was performed using either human recombinant IFNγ (Peprotech 300-02, 50ng/ml) and human recombinant TNFα (Peprotech 300-01A, 50ng/ml), human recombinant IFNα (Immunotools 11343516, 10ng/ml) and human recombinant IFNβ (Peprotech 300-02BC, 10ng/ml) or human recombinant TGFβ (Peprotech 100-21, 2ng/ml). Initially, cytokines were reconstituted in PBS containing 0.1% bovine serum albumin (BSA). Cells were treated with either the combination of IFNγ and TNFα for 24 or 48 hours, with the combination of IFNα and IFNβ for 48 hours or TGFβ for 48 hours. For the TGFβ condition, medium was refreshed with TGFβ after 24h.

### Lentivirus shRNAs

All shRNA targeting sequences were cloned into a pLV[shRNA]-mCherry:T2A:Puro-U6> vector purchased from Vectorbuilder or into DECIPHER pRSI9-U6-(sh)-UbiC-TagRFP-2A-Puro^29^. shRNA targeting sequences were selected based on previous literature^29^ and the RNAi consortium at the Broad Institute (https://portals.broadinstitute.org/gpp/public/). The shRNA target sequences used in this paper were: CCTAAGGTTAAGTCGCCCTCG (shCtrl#1), CAACAAGATGAAGAGCACCAA (shCtrl#2), CCGCAATTCCTTCCGCTACAA (shL28), GTCACGGAGGATAAGATCAAT (shP1#1), CGTCAACATTGGGAGCCTCAT (shP1#3), GTATTCCCGATCTGCCATGTA (shL6). HEK293T (Sigma-Aldrich, 12022001) cells were used for lentiviral packaging using a second (pMDL RRE and psPAX2) or third-generation (pVSV-G, pRSV-REV and pMDLg/RRE) packaging system. Tumor cells were infected with shRNA lentivirus containing supernatant at a MOI=0.5 and transduction efficiency was measured by flow cytometry 96 hours after infection. This resulted in a transduction efficiency of 80-100% of target cells determined by mCherry (%) expression.

### HLA class I surface staining

To assess HLA class I surface expression, tumor cells were stained for 20 minutes with BV605^TM^ anti HLA-A, B, C (clone W6/32, BioLegend 311432) and APC anti HLA-A2 (clone BB7.2, BioLegend 343308). Cell viability was measured using the LIVE/DEAD Fixable IR Dead Cell Stain Kit (Thermo Fisher L10199). Flow cytometric data were acquired using the BD FACSymphony^TM^ flow cytometer (BD Biosciences), gated on single and live cells. Data were analyzed with the FlowJo version 10.8.1 software (FlowJo LLC).

### HLA class I peptide complex recovery

Tumor cells were washed with PBS and treated for 2 minutes with ice cold citric acid buffer (0.13M citric acid, 0.061 M Na2HPO4, 0.15 M NaCl at pH 3). Cells were washed three times with ice cold PBS, resuspended in culture medium and kept at 37°C. At indicated time points, cells were stained with HLA-A, B, C (clone W6/32, BioLegend 311432) and APC anti HLA-A2 (clone BB7.2, BioLegend 343308) and HLA surface levels were measured on the BD FACSymphony^TM^ flow cytometer (BD Biosciences) using the MFI.

### Co culture experiments

MART-1 or NY-ESO-1 specific CD8^+^ T cells were thawed one day prior to the experiment in RPMI containing Penicillin - Streptomycin, 10% HS and DNase I (Sigma-Aldrich 10104159001, 1/1000) for 30 minutes. Cells were recovered overnight in RPMI supplemented with Penicillin - Streptomycin, 10% HS, IL-7 (5ng/ml) and IL-15 (5ng/ml). T cells and RP KD tumor cells were then co-cultured for 6 hours in the presence of GolgiPlug (BD Biosciences 555029) and GolgiStop (BD Biosciences 554724) at a T cell:target ratio of 1:1. After 6 hours, cells were stained with AF700 anti CD107a (clone H4A3, BD Biosciences 561340), BUV anti CD8 (clone SK1, BD Biosciences 612889) and the LIVE/DEAD Fixable IR Dead Cell Stain Kit. Subsequently, cells were fixed and permeabilized with the Foxp3 Transcription Factor Staining Buffer Set (eBioscience, 00-5523-00). Intracellular cytokine staining was performed using APC anti IFNγ (clone B27, BD Biosciences 554702) and FITC anti TNFα (clone MAb11, BD Biosciences 554512). Fixed cells were measured the following day using the BD FACSymphony^TM^ flow cytometer (BD Biosciences), gated on single, live CD8^+^ T cells. Data were analyzed with the FlowJo version 10.8.1 software (FlowJo LLC).

### Incucyte killing assay

MART-1 or NY-ESO-1 specific CD8^+^ T cells were thawed one day prior to the experiment as described above. To measure tumor cell death, tumor cells were stained with the Incucyte Caspase-3/7 Green Dye (Satorius 4440, 5µM). Subsequently, MART-1 or NY-ESO-1 specific CD8^+^ T cells were added at a T cell:target ratio of 1:1. As a negative control, tumor cells were cultured without T cells. After allowing cells to settle for 30 minutes, live-cell fluorescent images were captured every 2 hours with the Incucyte Live-Cell Analysis System (Satorius) for 48 hours in total. Of note, shL28 tumor cells were hypersensitive to cell contact induced apoptosis, which resulted in increased tumor death after 24h in both cell lines expressing shL28. Therefore, these lines were analyzed only until the 24h time point. T cell killing was assessed with the integrated Incucyte Base Analysis Software (Satorius) using dead tumor cell count. Importantly, each data timepoint was analyzed to the negative control to account for tumor death without T cells.

### ^35^S-methionine incorporation assay

Assessment of protein synthesis using ^35^S-methionine labelling was performed as described previously^61^. In short, tumor cells were labelled with 30 µCi/ml ^35^S-methionine label (Hartmann Analytic ARS0110) for 2 hours at 37°C. After washing cells with PBS, cells were lysed with lysis buffer (50 mM TrisHCl pH 7.5, 150 mM NaCl, 1% Tween-20, 0.5% NP-40, protease inhibitor cocktail (Roche 5892791001) and phosphatase inhibitor cocktail (Sigma P5726). Protein was precipitated onto Whatmann filter paper using 25% trichloroacetic acid (TCA). After washing with 70% ethanol and acetone, scintillation was measured with the liquid scintillation counter (Perkin Elmer). For data analysis, radioactive counts were normalized to total protein amounts.

### Immunoblotting

Tumor cells were washed in PBS and pellets were lysed for 20 minutes on ice in lysis buffer (100mM Hepes pH 7.5, 300mM NaCl, 10mM EGTA, 3mM MgCl_2_, 2% glycerol, 2% triton X-100 (Sigma), protease inhibitor cocktail (Roche 5892791001) and phosphatase inhibitor cocktail (Sigma P5726). Protein amounts were quantified with the Pierce™ BCA Protein Assay Kit (Thermo Scientific 23227) and equal protein amounts were separated using ready-to-use 4–20% Mini-PROTEAN® TGX Stain-Free™ Protein Gels (Biorad, 4568096) and transferred onto 0.2 μm pore nitrocellulose membranes (PALL Life Sciences 66485).

Subsequently, dry membranes were blocked for 1 hour at room temperature using Intercept (TBS) Blocking Buffer (LI-COR, 927-66003). After blocking, membranes were incubated with primary antibodies overnight at 4°C followed by incubation with secondary antibodies IRDye® 800CW Goat anti-Rabbit IgG (LI-COR 926-32211, 1: 10000) and IRDye® 680RD Goat anti-Mouse IgG (LI-COR, 926-68070, 1:10000) for 1 hour at room temperature. Data were acquired with the Odyssey CLx (LI-COR) and analysed using the software ImageStudioLite version 5.2.5 (LI-COR) or Empiria Studio version 1.3.0.83 (LI-COR).

Primary antibodies used were: α-Tubulin (DM1A, Cell Signaling Technology 3873, 1:1000), HSP 90α/β (Santa Cruz Animal Health sc-13119, 1:1000), RPL28 (Abcam ab138125, 1:1000), RPLP1 (LifeSpan Biosciences LS-C198116-100, 1:1000), RPLP2 (Abcam ab154958, 1:1000), HLA Class 1 ABC (Abcam, ab70328, 1:1000), MART-1/Melan-A (Thermo Fisher MS-716-P, 1:1000), NY-ESO-1 (Thermo Scientific PA5-116201, 1:500), Phospho-eIF2α (Ser51) (Cell Signaling Technology 9721, 1:1000), RPL7 (Thermo Scientific PA5-36571, 1:1000), RPL22 (Novus Biologicals NBP1-98446, 1:1000) and HA-Tag (Cell Signaling Technology 3724, 1:1000).

### RNA isolation and RT-qPCR

RNA was isolated with the ISOLATE II RNA Mini Kit (BioCat BIO-52072-BL) according to the manufacturer’s protocol. Isolated RNA was resuspended in nuclease-free water and quantified using the Nanodrop (Thermo Scientific). Subsequently, High-Capacity cDNA Reverse Transcription Kit (Thermo Scientific 4368814) was used for reverse transcription and SYBR^TM^ Green PCR Master Mix (Thermo Fisher 4367659) was used for qPCR. Gene expression levels were analysed in three technical replicates by using the comparative CT method and data were normalised to the housekeeping gene *ACTB.* The used primer sequences for qPCR were: *ACTB* forward (5’-CCTGGCACCCAGCACAAT-3’), *ACTB* reverse (5’-GGGCCGGACTCGTCATACT-3’), *RPL28* forward (5’-GTGCCTCAGTTTCCCATATGTA-3’), *RPL28* reverse (5’-TACTGGACTAAGAGCTGGAGAG-3’), *RPLP1* forward (5’-CCTATACAACCCTGCCTAAGAAC-3’), *RPLP1* reverse (5’-CCCAAACATTTGGTGAGACATTAC-3’), *HLA-A* forward (5’-AAGAGTTGTTCCTGCCCTTC-3’), *HLA-A* reverse (5’-CCTCCTCACATTATGCCTACAC-3’), *HLA-B* forward (5’-GACACTGAGCTTGTGGAGAC-3’), *HLA-B* reverse (5’-GGCATGTGTATCTCTGCTCTT-3’), *HLA-C* forward (5’-AATGTGAGGAGGTGGAGAGA-3’), *HLA-C* reverse (5’-CCTCTCTGGAACAGGAAAGATG-3’).

### Polysome fractionation (protein precipitation and RNA extraction)

Tumor cells were lysed in ice cold polysome lysis buffer (10x gradient buffer (1.1M potassium acetate, 0.2M magnesium acetate, 0.1M HEPES pH 7.6), 100mM KCl, 10mM MgCl_2_, 0.1% NP-40, 154.25mg/ml DTT, 100 μg/ml cycloheximide (Sigma-Aldrich C7698), RNaseOUT™ Recombinant Ribonuclease Inhibitor (Thermo Scientific 10777019) and combined Halt^TM^ Protease and Phosphatase Inhibitor EDTA-free (Thermo Scientific 78443)). Lysates were passed through a needle (27G) and cleared by centrifugation at 14.000 x g for 10 minutes. Subsequently, 50ul of cleared supernatant was collected to measure the total protein or RNA amount while the rest was loaded onto a 10%-60% sucrose gradient and centrifuged under vacuum at 38.000 rpm for 2 hours at 4°C. Subsequently, monosome-containing fractions 6-8 (750ul per fraction) and polysome-containing fractions 9-14 (750ul per fraction) were collected and protein was extracted using a standard TCA precipitation protocol^62^.

For RNA extraction from sucrose gradients, each fraction (fraction 1-14) was collected separately and incubated with 20mg/ml Proteinase K (Thermo Scientific 25530049, 1:100), 0.5M EDTA and 20% SDS for 30 minutes at 42°C. Afterwards, samples were mixed with phenol:chloroform and centrifuged for 8 minutes at 14.000 x g. Subsequently, RNA was extracted with 100% EtOH and sodium acetate and centrifuged for 30 minutes at 14.000 x g. Samples were then washed with 80% EtOH and centrifuged for 10 minutes at 14.000 x g.

Finally, RNA pellets were reconstituted in nuclease-free water, incubated for 5 minutes at 60°C and stored at −80°C.

### Protein quantification with LC-MS

#### Quantification of ribosomal fractions

Dry protein pellets were dissolved in 100mM TrisHCL (pH 8.5, Sigma T1503) to determine the protein concentration using the Bradford protein assay (Thermo Scientific 23200). Subsequently, proteins were reduced and alkylated with dithiothreitol (Sigma 43815) and iodoacetamide (Sigma I6125) and digested twice with trypsin (Promega VA9000) at 37°C. The digests were dried, reconstructed with 5% formic acid (Honeywell|FlukaTM 94318) and desalted with C18 StageTips (Thermo Scientific 87781). Before proceeding with mass spectrometry, the digests were reconstructed with 2% formic acid. All samples were analyzed using a nanoLC-MS/MS system consisting of an Orbitrap Exploris 480 mass spectrometer coupled to an EASY-NLC 1200 system (Thermo Scientific). The proteins were separated on an analytical column (ReproSil-Pur 120 C18-AQ, 2.4 μm, 75 μm × 500 mm, packed in-house) using a linear gradient of solvent A (0.1% formic acid/water, Fischer Chemical LS122) and solvent B (0.1% formic acid/80% acetonitrile, Fischer Chemical LS118), with a flow rate of 250 nl/minutes in a 90-minutes gradient. The gradient contained a 70-minutes linear increase from 6% to 30% solvent B, followed by a 20-minutes wash at 90% solvent B.

#### Total protein quantification

For protein digestion, frozen cell pellets were lysed in boiling Guanidine (GuHCl) lysis buffer as described by Jersie-Christensen et al^63^. Protein concentration was determined with a Pierce Coomassie (Bradford) Protein Assay Kit (Thermo Scientific), according to the manufacturer’s instructions. After dilution to 2M GuHCl, aliquots of protein were digested twice (overnight and 4h) with trypsin (Sigma-Aldrich) at 37°C, enzyme/substrate ratio 1:75. Digestion was quenched by the addition of FA (final concentration 1%), after which the peptides were loaded on the Evotip Pure™ (Evosep). LC-MS/MS analysis was carried out using an Evosep One LC system connected to an Orbitrap Exploris 480 Mass Spectrometer (Thermo Fisher Scientific). Peptides were separated using the pre-programmed gradient (Extended method, 88 min gradient) on an EV1137 (Evosep) column with an EV1086 (Evosep) emitter. The Exploris 480 was run in data-independent acquisition (DIA) mode, with full MS resolution set to 120,000 at m/z 200, MS1 mass range was set from 350-1400, normalized AGC target was 300% and maximum IT was 45ms. DIA was performed on precursors from 400-1000 in 48 windows of 13.5 Da with an overlap of 1 Da. Resolution was set to 30,000 and normalized CE was 27.

#### Data analysis

For both runs, raw data were analyzed by DIA-NN (version 1.8)^64^ without a spectral library and with “Deep learning” option enabled. The Swissprot human database (20,395 entries, release 2021_04 for ribosomal fractions MS; 20.398 entries, release 2022_08 for P1-KD MS) was added for the library-free search. The Quantification strategy was set to Robust LC (high accuracy) and MBR option was enabled. The other settings were kept at the default values. For ribosomal fractions MS, the protein groups report from DIA-NN was used for downstream analysis in R, whereas, for the P1-KD MS, protein group abundances were extracted from the DIA-NN result files, imported into Perseus (version 2.0.7.0)^65^ and were filtered for presence in at least 3 out of 4 replicates in at least one condition. Further analyses were performed in R as follows. For the ribosome-specific analyses of sub-polysome and polysome fractions, samples were first clustered into different fractions and for each independent cluster, RP abundances were normalized using a variance stabilizing normalization (vsn)^66^, using solely the abundances of detected RPs. For total protein fraction, we performed the *vsn* in the presence of all detected proteins. For ratio analysis, normalized abundances of each RP were replicate-matched and divided between different fractions, creating the replicate-specific polysome/total, subpoly/total and sub-poly/poly ratios. Statistical significance of differential protein abundance and ratio values was obtained using a two-sided T-test and Benjamini-Hochberg p-value adjustment. Gene Set Enrichment Analyses and its visualization were performed using the clusterProfiler package^67^, for which the genesets were taken from MSigDB v7.0 and gene-ranking metric was selected as log2FC x min(-log10(adj.p-val), 3).

### Phosphoproteomics

#### Phosphopeptide enrichment

Phosphorylated peptides were enriched from 10uG total peptides using a High-Select Fe-NTA Phosphopeptide Enrichment Kit (Thermo Scientific A32992), according to the manufacturer’s instructions, with the exception that the resin was split into five aliquots, with each aliquot subsequently loaded onto a StageTip, C8 (Thermo Scientific). Centrifugation steps were performed at 1000 x g for 5 minutes and dried eluates were reconstituted in 15 μL of 2% formic acid.

#### Quantification of phosphopeptides

Prior to mass spectrometry analysis, the peptides were reconstituted in 2% formic acid. Peptide mixtures were analyzed by nanoLC-MS/MS on an Orbitrap Exploris 480 Mass Spectrometer equipped with an EASY-NLC 1200 system (Thermo Scientific). Samples were directly loaded onto the analytical column (ReproSil-Pur 120 C18-AQ, 2.4 μm, 75 μm × 500 mm, packed in-house). Solvent A was 0.1% formic acid/water and solvent B was 0.1% formic acid/80% acetonitrile. For the phosphoproteome, samples were eluted from the analytical column at a constant flow of 250 nl/min in a 70-min gradient containing a linear increase from 7% to 30% solvent B, followed by a 15-min wash at 90% solvent B.

#### Data analysis phosphoproteomics

For phosphoproteome, data (RAW files) were analyzed by MaxQuant (version 2.0.1.0) ^68^ using standard settings. MS/MS data were searched using the Swissprot human database (20,395 entries, release 2021_04) complemented with a list of common contaminants and concatenated with the reversed version of all sequences. Trypsin/P was chosen as cleavage specificity allowing two missed cleavages. Carbamidomethylation (C) was set as a fixed modification, while oxidation (M) and phosphorylation(S,T,Y) were used as variable modifications. LFQ intensities were Log2-transformed in Perseus (version 1.6.15.0) ^65^, after which the phosphosites were filtered for at least two valid values (out of 3 total) in at least one condition. The values were normalized by subtracting the median per sample. Missing values were replaced by imputation based on a normal distribution using a width of 0.3 and a downshift of 1.8. Differentially regulated phosphosites were determined using a t-test (threshold:–log(p-value) ≥ 1.3 and [x-y] ≥ 1 | [x-y] ≤ −1).

### Sample preparation and HA pulldown

For the HA pulldown, eL22-HA and HA-P1 tumor cells were grown in 4×15cm dishes per condition. Cytokine-treated HA tagged tumor cells were treated with 50ng/ml IFNγ and TNFα for 24 hours. As described previously^61^, cells were treated with 200 μg/ml cycloheximide for 3-5 minutes at 37 °C and cell pellets were washed with ice-cold PBS containing 200 μg/ml cycloheximide. Subsequently, pellets were lysed for 30 minutes in ice-cold lysis buffer (20 mM Tris HCl pH 7.4, 10 mM MgCl2, 150 mM KCl, 1% NP-40, 200 µg/mL cycloheximide and 1x EDTA-free proteinase inhibitor cocktail (Sigma 5892791001)). Lysates were centrifuged at 12.000 x g for 20 minutes at 4 °C and the supernatants were collected for RNA digestion using RNaseI (Thermo Scientific AM2294) for 40 minutes at 25 °C. RNA digestion was stopped using SUPERase (Thermo Scientific AM2694). Digested lysates were incubated with Pierce™ Control Agarose Matrix (Thermo Scientific 26150) for 15 minutes at 4°C followed by incubation with AntiHA.11 Epitope Tag Affinity Matrix (BioLegend 900801) for 4 hours at 4°C under constant rotation. Supsequely, tagged ribosomes were eluted with 1mg/ml HA peptide (Thermo Scientific 26184) for 15 minutes at 30°C. Finally, ribosome protected fragments (RPFs) were purified using miRNeasy minikit (Qiagen) according to manufacturer’s instructions and RNA concentrations were measured with the Agilent 2100 bioanalyzer using a Small RNA Chip (Agilent 5067-1548).

### Ribosome profiling

Ribosome profiling was performed in technical duplicates on purified RPFs of cytokine-treated HA tagged Mel624 tumor cells as described previously^69^. Briefly, RPFs were run on a 15% TBE-urea polyacrylamide gel and size-selected between 26nt and 32nt using RNA markers. RNA was extracted using RNA gel extraction buffer (300 mM NaOAc pH 5.5, 1.0 mM EDTA, 0.25% SDS) and precipitation for 30 minutes on dry ice. After precipitation, samples were incubated overnight at 25°C. End dephosphorylation was performed using T4 polynucleotide kinase (PNK) (NEB M0201) for 1 hour at 37°C and ligation was performed using the T4 RNA ligase I (NEB M0204) for 2 hours at 25°C. Reverse transcription was performed using the SuperScript III reverse transcriptase (Thermo Scientific 18080051) using the oCJ485 RT primer for 30 minutes at 50°C. cDNA was then size-selected at 110bp using a 10% TBE-urea polyacrylamide gel. DNA was extracted using DNA gel extraction buffer (300 mM NaCl, 10 mM Tris pH 8, 1 mM EDTA) and precipitation for 30 minutes on dry ice. After precipitation, samples were incubated overnight at 25°C. Circular ligation was performed using CircLigase II ssDNA Ligase (Epicentre CL4111K) for 1,5 hours at 60°C. cDNA amplification was performed for 15 cycles with the Phusion High-Fidelty DNA polymerase (Thermo Scienfific F530) and primers containing different indexes for sequencing. The amplification product was size-selected at 180bp using a 8% TBE polyacrylamide gel. Samples were measured with the Agilent 2100 bioanalyzer using a DNA chip (Agilent 5067-1506) to assure high quality followed by molarity quantification. Sequencing was performed on the Illumina HiSeq2500.

### Ribo-seq data analysis

Sequencing data from Ribo-seq experiments was first adapter-trimmed using the cutadapt tool^70^, then, after cleaning rRNA & tRNA fragments, remaining reads were mapped to hg38 genome using the STAR aligner^71^. QC plot (RPF length and periodicity) generation and translated ORF predictions were performed with RiboCode^72^ using the gencode v34 annotation. Differential analysis of ORF abundances was performed with the DESEQ2 package^73^ in the R environment, where ORF-mapping reads were quantified with the Rsubread package^74^. Gene Set Enrichment Analyses were performed with the clusterProfiler^67^ package using the log2FC x min(-log10(p-value),5) measure when ranking genes. For the codon level analysis of ribosome occupancy (RO) around transmembrane domain (TMD) ends, TMD coordinates were retrieved from ENSEMBL96 annotation.

Focusing only on primary transcripts (based on APPRIS annotation), sample-specific ribosome P-site occupancies (position 12-14 within RPF) were counted at each codon distance from the reference coordinates using a custom python script. For the figure, distance-specific RO changes were calculated as logFC between two groups, based on group-specific replicate-averaged normalized read count sum across all given reference coordinates, and plotted using a custom R script.

### TCGA data analysis

Gene expression (HTSeq FPKM) values for all TCGA cohorts were downloaded via UCSC Xena project^41^. For each cohort, sample-specific geneset signature scores were computed as normalized enrichment scores using the ssGSEA method. For each sample, relative RP expression values were obtained with a z-score transformation of raw FPKM values of all ribosomal proteins, excluding the ribosomal protein pseudogenes. All data handling and plotting were performed with a custom R script.

### Data and Software Availability

The proteomics data have been deposited at the PRIDE repository under accession number PXD041688. The RiboSeq data have been stored at Gene Expression Omnibus (GEO) with the ID GSE230060.

## Author Contributions

AD, PK, and WJF designed and conceived the study. Data were generated and analyzed by AD, FA, YM, KH, MK, AG, JS, SR, LH, and RvdK. Computational analyses were carried out by FA and OI. Manuscript was written and edited by AD, JWY, PK, and WJF. All authors provided feedback on the analyses, edited and approved the manuscript.

---

**Supp Figure 1:** Cytokine exposure induces an “alert-state” in tumor cells **A.** IFNγ and TNFα treatment up-regulates HLA I surface levels in Mel624 and M026 tumor cells. HLA levels were measured by flow cytometry using MFI. n=3 independent experiments per cell line each assessed in triplicates represented as mean ± SEM. p values were calculated using a two-tailed t-test. **B.** IFNγ and TNFα treatment increases immunoproteasome subunits (IP; PSMB8, PSMB9, PSMB10) compared to standard proteasome subunits (SP; PSMB5, PSMB6, PSMB7) in Mel624 and M026 tumor cells. n=3 independent experiments per cell line represented as mean. **C.** IFNα and IFNβ treatment up-regulates HLA I surface levels in tumor cells. HLA levels were measured by flow cytometry using MFI. n=3 independent experiments per cell line each assessed in triplicates represented as mean ± SEM. p values were calculated using a two-tailed t-test. **D.** IFNα and IFNβ treatment increases immunoproteasome subunits (IP; PSMB8, PSMB9, PSMB10) compared to standard proteasome subunits (SP; PSMB5, PSMB6, PSMB7). n=3 independent experiments per cell line represented as mean. **E-G.** Western blot analysis confirms increased polysomal association of P1 upon IFNγ and TNFα stimulation in SK-OV-3 (G), HCT 116 (H) and hTERT-RPE1 (I) cells. uL30 was used as control RP for data normalization. n=1 experiment per cell line.

**Supp Figure 2:** P1 KD results in decreased protein synthesis and cell proliferation **A.** Transduction efficiency was measured by flow cytometry using mCherry (%) expression, gated on live tumor cells. n=3 independent experiments per cell line each assessed in triplicates represented as mean ± SEM. **B.** Western blot analysis confirming efficient RP KD. α-tubulin was used as loading control. n=1 experiment per cell line. **C.** RT-qPCR analysis confirming efficient RP KD, using *ACTB* as a reference. n=3 technical replicates represented as mean ± SD. p values were calculated using a two-tailed t-test. **D.** Western blot demonstrating a decrease in HLA I protein levels in shP1 compared to control M026 tumor cells. HSP90 was used as loading control. n=1 independent experiment. **E.** RT-qPCR showing no change in HLA mRNA levels (*HLA-A, HLA-B, HLA-C*) following P1 KD, using *ACTB* as a reference. n=2 independent experiments each assessed in technical triplicates. Data are represented as mean ± SEM and p values were determined using a two-tailed t-test.

**Supp Figure 3:** The ASR translates mRNAs important for immunosurveillance **A.** Schematic representation of ribosome profiling experimental set up. **B.** Western blot confirming the expression of tagged HA-eL22 and P1-HA (input: total, pre-clearance (pre), supernatant (super)) and isolation of eL22- and P1-containing ribosomes (Immunoprecipitation (IP)) in Mel624 tumor cells. Co-IP of uL30 was used as a control for pulldown of intact ribosomes and HSP90 was used as loading control. n=1 replicate per condition. **C.** Ribo-seq quality control plots showing the length distribution and periodicity of sequenced ribosome protected fragments in our Ribo-seq experiments. **D.** Pie chart showing the distribution of predicted ORF classes, performed by RiboCode separately on P1-PD and L22-PD Ribo-seq data allowing alternative start codons. **E.** Predicted ORF class distribution, performed by RiboCode jointly on all Ribo-seq samples, allowing only ATG start codon. **F.** PCA with ORF-mapping read quantifications for which Ribo-seq reads were quantified by the subread package on ORFs. **G.** Proteomics analysis of the M026 cell line reveals down-regulation of HLA and other APP components upon P1 KD. n=3 independent experiments and p values were calculated using a two-tailed t-test. **H.** Ribosome profiling-based GSEA of Mel624 cells shows that P1-containing ribosomes have increased translation rate of gene sets associated with antigen presentation. n=2 biological replicates and p values were calculated using the *clusterProfiler* package. Data demonstrate top 10 genesets and APP genesets.

**Supp Figure 4:** The ASR regulates cytokine-mediated processes **A.** KD of P1 decreases HLA I surface levels in M026 cells. HLA levels were assessed by flow cytometry using the MFI. n=3 independent experiments each assessed in triplicates. Data are represented as mean ± SEM and p values were determined using a two-tailed t test. **B.** shP1 M026 tumor cells showing slower recovery kinetics of HLA I after acid wash compared to the control. n=3 independent experiments. Data are represented as mean ± SEM. **C.** Exposure to IFNγ and TNFα results in attenuated basal HLA I surface levels in shP1 M026 tumor cells. n=3 biological replicates represented as mean ± SEM. p values were calculated using a two-tailed t-test. **D.** MART-1-specific CD8^+^ T cell recognition is decreased in shP1 compared to control M026 tumor cells. n=3 independent experiments each assessed in triplicates. Data are represented as mean ± SEM and p values relative to shCtrl were determined using a two-tailed t-test. **E.** NY-ESO-1-specific CD8^+^ T cell recognition is decreased in shP1 compared to control M026 tumor cells. n=3 independent experiments each assessed in triplicates. Data are represented as mean ± SEM and p values relative to shCtrl were determined using a two-tailed t-test. **F.** MART-1 specific T cell killing is not suppressed in shP1 M026 tumor cells. n=3 independent experiments per cell line, each assessed in triplicates. Data represent mean and error ± SEM and p values were calculated with a two-tailed t-test. **G.** NY-ESO-1 specific T cell killing is suppressed in shP1 M026 tumor cells. n=3 independent experiments per cell line, each assessed in triplicates. Data represent mean and error ± SEM and p values were calculated with a two-tailed t-test.

**Supp Figure 5:** P-stalk levels correlate with T cell response across diverse tissues **A.** Western blot showing unaltered MART-1 and NY-ESO-1 source protein levels upon P1 KD in Mel624 and M026 cells. α-tubulin was used as loading control. n=2 independent experiments represented as mean ± SEM and p values were calculated with a two-tailed t-test. **B.** ^35^S-methionine incorporation assay showing decreased protein synthesis upon P1 KD in Mel624 and M026 cells. Cycloheximide (CHX) was used as a control to inhibit protein synthesis. n=3 technical replicates represented as mean ± SD. p values were calculated using a two-tailed t-test. **C.** CellTrace violet cell proliferation assay showing decreased cell proliferation upon P1 KD in M026 and Mel624 cells. Proliferation was assessed by flow cytometry using the MFI. n=3 biological replicates represented as mean ± SEM. p values were calculated using a two-tailed t-test. **D.** ^35^S-methionine incorporation assay showing inhibited protein synthesis in Mel624 and M026 cells following eL6 KD similar to P1 KD. n=3 technical replicates represented as mean ± SD. p values were determined using a two-tailed t-test. **E.** Translation inhibition caused by KD of eL6 shows minor effects on HLA I surface levels in Mel624 and M026 tumor cells. n=3 biological replicates represented as mean ± SEM. p values were calculated using a two-tailed t-test. **F.** Western blot analysis showing no increase of elF2α phosphorylation following P1 KD, with or without IFNγ and TNFα. HSP90 was used as loading control. n=1 experiment per cell line. **G.** Gene expression analysis in the TCGA-SKCM melanoma cohort reveals a significant correlation between P1 and P2 relative expression levels (z-score of FPKM values within all RPs) and effector CD8 T cell infiltration. P values were calculated using Spearman’s rank correlation. n=474 **H.** TCGA-SKCM cohort gene expression analysis reveals a significant correlation between P1 and P2 relative expression levels and IFNγ response signature. P values were calculated using Spearman’s rank correlation. n=474

