## Supplementary figures and images for "An “alert state” ribosome population acts as a master regulator of cytokine-mediated processes"

### Supp. Fig. 1

Supp. Figure 1

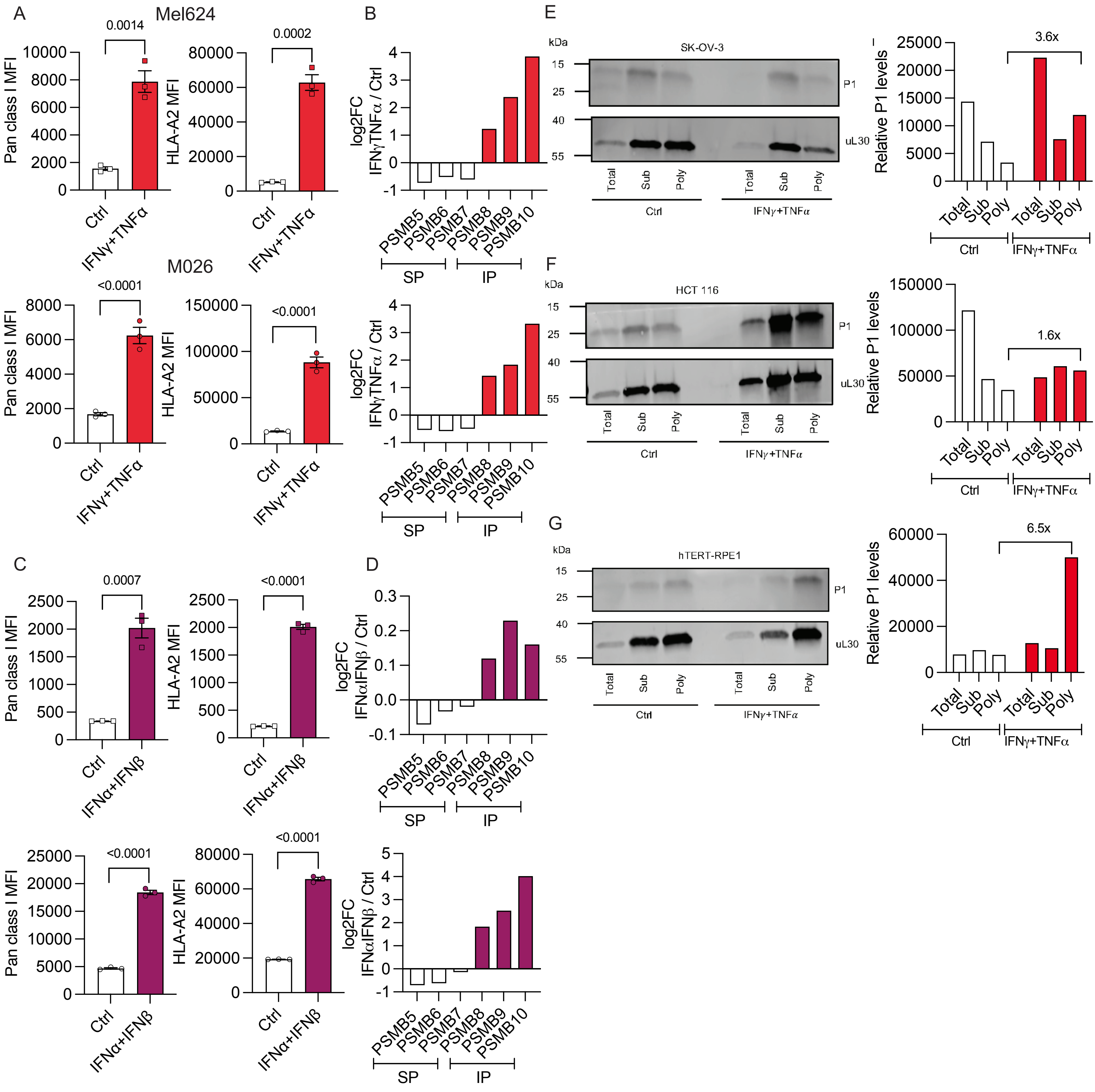

### Supp. Fig. 2

#### Supp. Figure 2

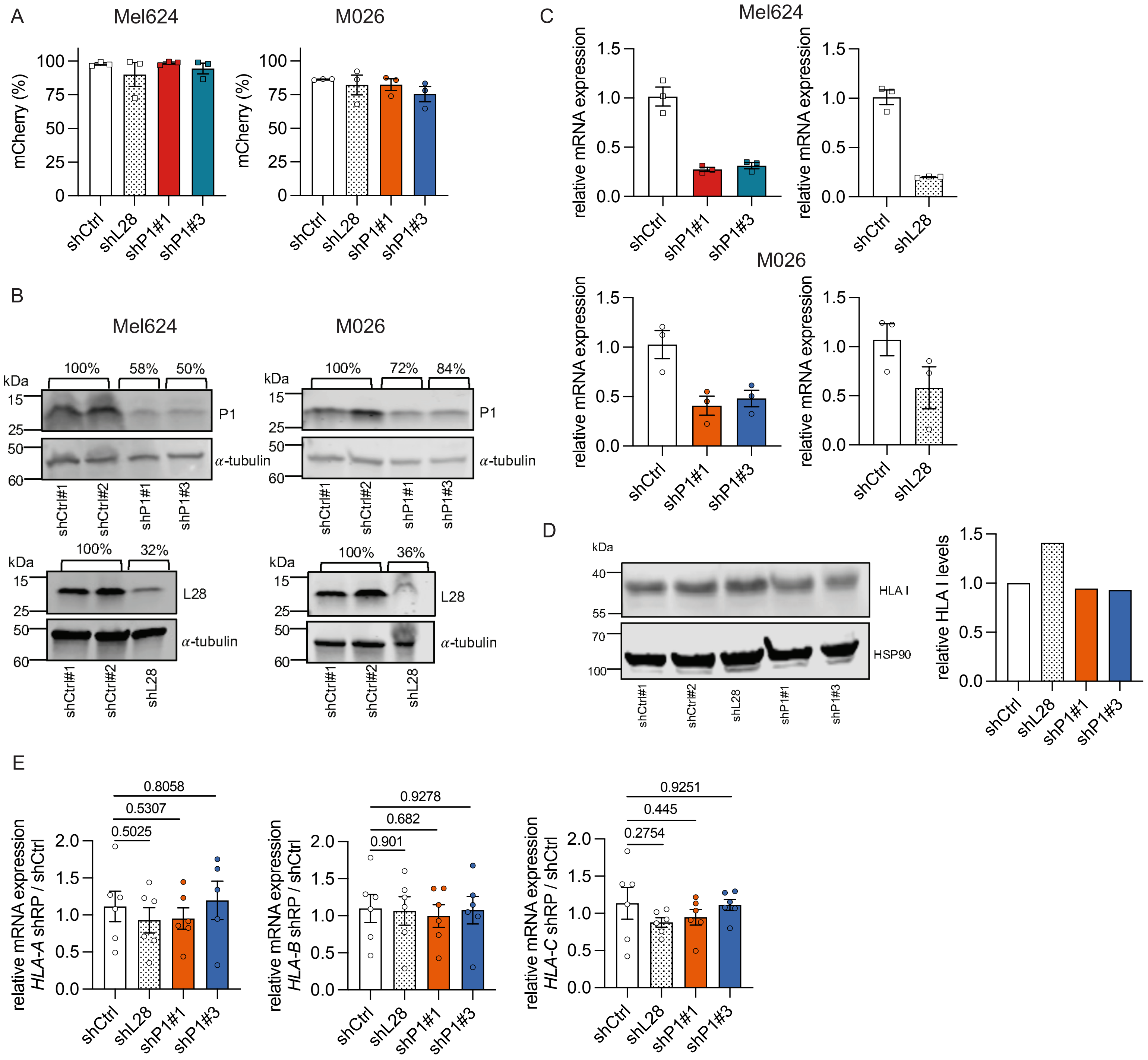

### Supp. Fig. 3

Supp. Figure 3

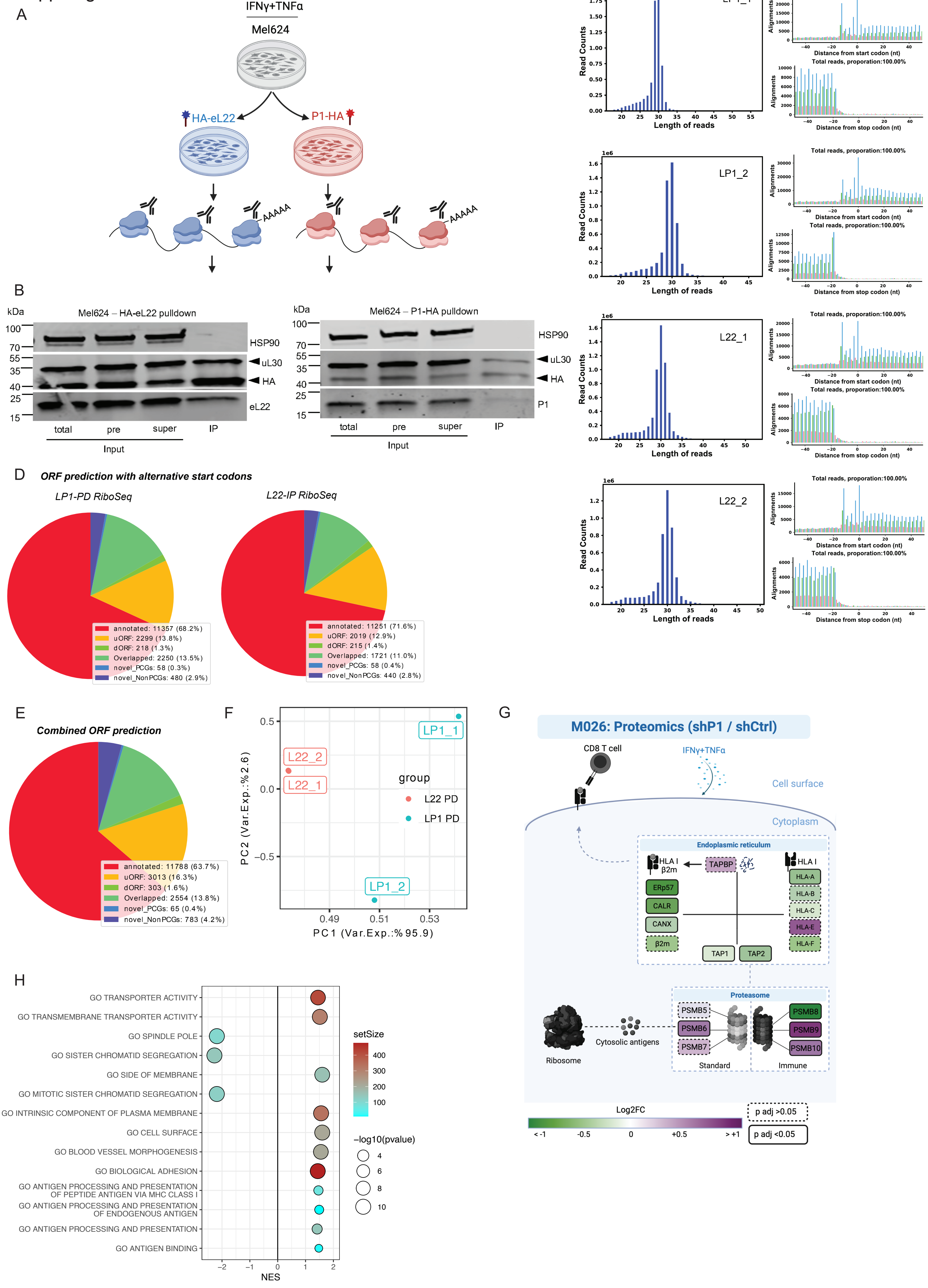

### Supp. Fig. 4

Supp. Figure 4

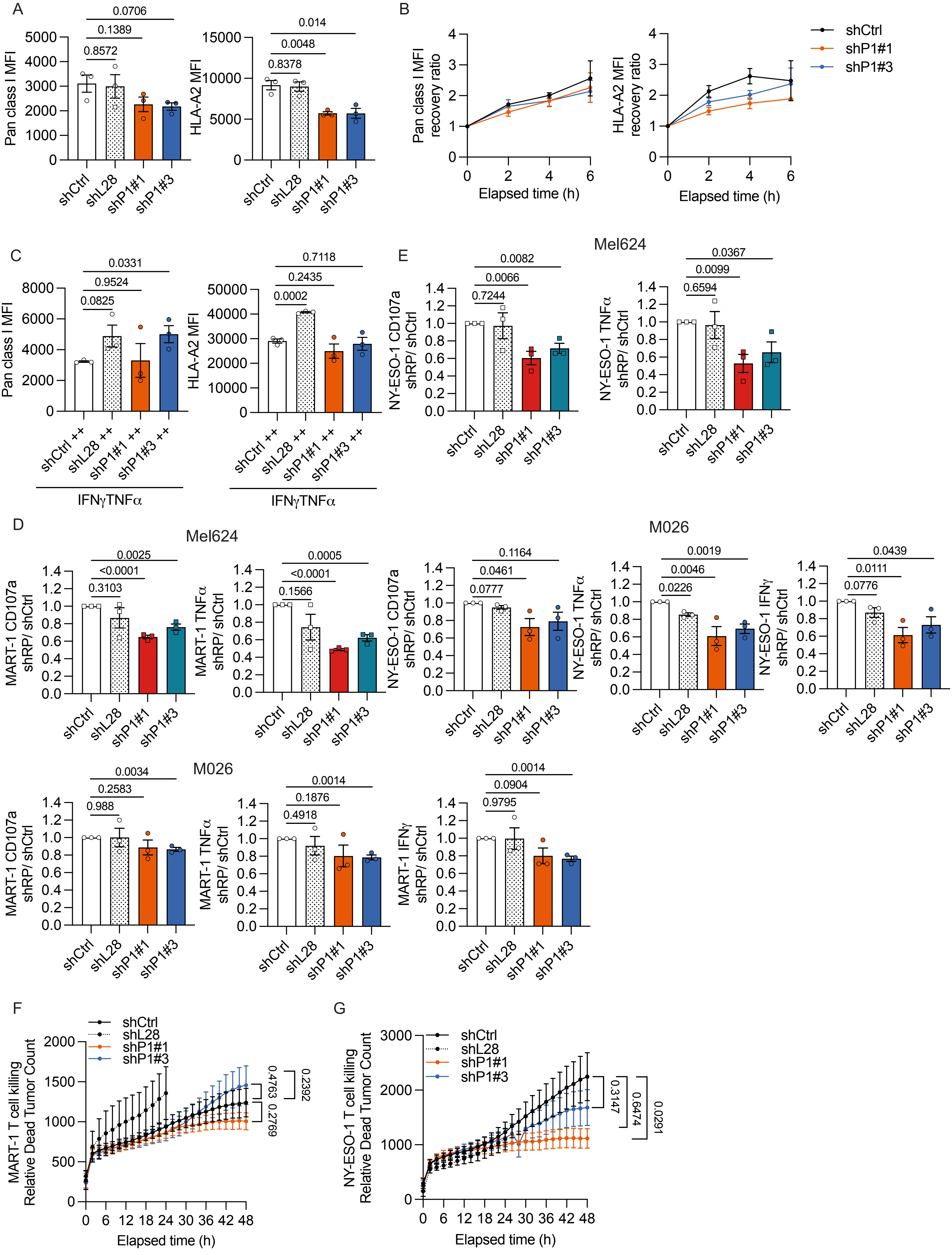

### Supp. Fig. 5

Supp. Figure 5

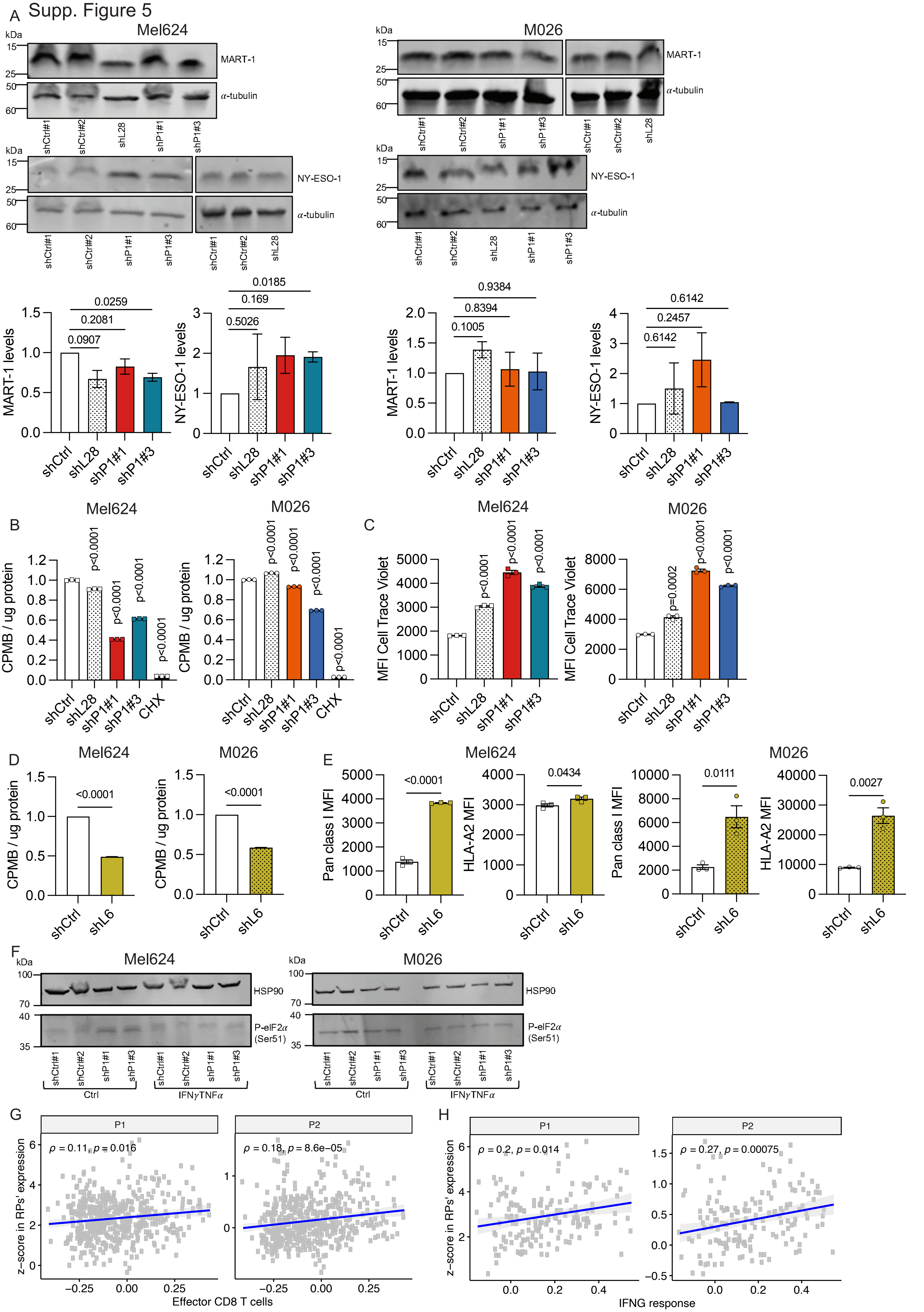
